# The trade-off between growth and risk in Kelly’s gambling and beyond

**DOI:** 10.1101/2023.11.07.566039

**Authors:** S. Cavallero, A. Rousselot, R. Pugatch, L. Dinis, D. Lacoste

## Abstract

We study a generalization of Kelly’s horse model to situations where gambling on horses other than the winning horse does not lead to a complete loss of the investment. In such a case, the odds matrix is non-diagonal, a case which is of special interest for biological applications. We derive a trade-off for this model between the mean growth rate and the volatility as a proxy for risk. We show that this trade-off is related to a game-theoretic formulation of this problem developed previously. Since the effect of fluctuations around the average growth rate is asymmetric, we also study how the risk-growth trade-off is modified when risk is evaluated more accurately by the probability of the gamble’s ruin.

## Introduction

In his seminal work from 1948, Shannon founded information theory. A pivotal contribution of Shannon’s theory was an existence proof he provided for a code that can allow signals to pass through a noisy channel with a negligible loss of information as long as the rate is smaller than the channel capacity. This was a big surprise back in the day since the belief up to that point was that noise monotonically reduces the information rate. Hence, from its outset, information theory involved coding. Enter J. R. Kelly. Shannon’s friend and colleague. In 1956, Kelly found a surprising example that hinted on a deep connection between information theory and gambling in horse races. With Shannon’s support, he published a paper, which is by now well-known and highly influential, titled “A new interpretation of information rate” [1]. In his article, Kelly showed, surprisingly, that the channel capacity also appears in the context of repeated bet-hedging on stationary horse races as the growth rate of the bettor’s capital.

Briefly, a horse race is said to be stationary if both odds and winning probabilities for each horse are constant in time. It can be shown that if one is trying to optimize the expected capital, one has to bet all the capital on the horse with the maximal distribution of winning. However, this will lead to the complete ruin of all the bettor’s capital in the long run. Kelly offered an alternative optimization criterion, which is by no means unique, namely, to optimize the growth rate of the capital. It then proceeded to show that in scenarios where partial information regarding the winning probability is provided by an insider (“side information”), the optimal strategy using his criterion can be calculated; it depends on the conditional probability of winning each horse and leads to a growth rate that is equal to the channel capacity of the insider-bettor information channel (where the information is the uncertain identity of the winning horse). As mentioned, there is no apparent coding involved in the scheme. Much later, Cover et. al. have devised a coding scheme similar to arithmetic coding that uses two identical bettors to code and decode messages.

Over the years, Kelly’s idea grew into a whole branch of information theory [2] with theoretical and practical implications for portfolio management [3] and more generally for investment strategies in finance [4]. Kelly’s model can also be formulated as a non-linear control problem with fruitful implications in finance and applied mathematics [5].

In the context of biology, Kelly’s work is important because it offers a way to connect information and fitness [6, 7], a central question in evolutionary biology. This connection is well illustrated with the notion of bet-hedging. Bet-hedging is a strategy that spreads the bets on phenotypes used by a population of individuals in which individuals accept a reduction of their short-term reproductive success, in exchange for longer-term risk reduction for the population [8–10]. Such a strategy is employed for instance by cells to cope with antibiotics [11], by phages to optimize their infection strategy of bacteria [12], or by plants to cope with a fluctuating climate [13, 14]. All these three examples involve a dormant state which protects individuals from harsh conditions in the environment, while preserving the biodiversity in a seed bank [15].

In the context of gambling, Kelly’s strategy is known to be risky, and for this reason most gamblers use fractional Kelly’s strategies, with reduced risk and growth rate [3]. This observation hints at a trade-off between the risk the gambler is ready to take and the average long-term growth rate of his capital, which is known in finance under the name of risk-return trade-off. In previous work, we have studied this trade-off in a version of Kelly model with a risk constraint [16]. In subsequent work, we found a similar trade-off in the context of a biological population with phenotypic switching in a fluctuating environment [17] by building on a piece-wise Markov model introduced earlier [18]. We have also proposed an adaptive version of Kelly’s gambling based on Bayesian inference [19].

In this paper, we study the risk-return trade-off for a generalization of Kelly’s model to situations where the gambling on horses other than the winning horse does not lead to a complete loss of the investment [6, 20, 21]. We develop a game-theoretic formulation of this problem building on Ref. [22]. Finally, we explore an alternate measure of risk, based on the probability of bankruptcy of the gambler, rather than the volatility, and we discuss some consequences of these ideas for biology or ecology.

### 1 Definition of Kelly’s model

Let us recall the basic elements of Kelly’s horse race model [1]. A race involves *M* horses, and is described by a normalized vector of winning probabilities **p**, an inverse-odds vector **r** and a vector of bets which defines the gambler strategy **b**. The latter corresponds to a specific allocation of the gambler’s capital on the *M* horses: if we denote by *C*_*t*_ the gambler’s capital at time *t*, the amount of capital invested on horse *x* reads b_*x*_*C*_*t*_. We further assume that, after each race, the gambler invests his whole capital, i.e.,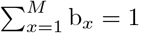 = 1, always betting a non-zero amount on all horses, i.e., *∀x ∈* [1, *M*] : b_*x*_ */*= 0. The inverse-odds vector **r** is set by the bookmaker. When Σ_*x*_ r_*x*_ = 1, the odds are *fair*, when Σ_*x*_ r_*x*_ *>* 1 the odds are *unfair* and when Σ_*x*_ r_*x*_ *<* 1 the odds are *superfair*.

The evolution of the gambler’s capital after one race reads:

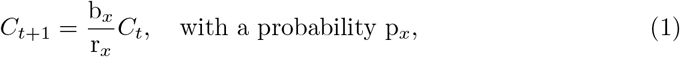

which implies that the log of the capital, log-cap(*t*) *≡* log *C*_*t*_, evolves additively:

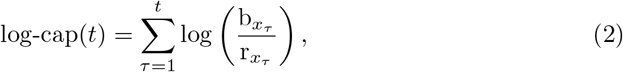

where *x*_*τ*_ denotes the index of the winner of the *τ* -th race and we assumed log-cap(0) = 0 (i.e. *C*_0_ = 1). Since the races are assumed to be independent, the terms log 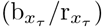 in (2) are independent and identically distributed, and we can use the weak law of the large numbers :

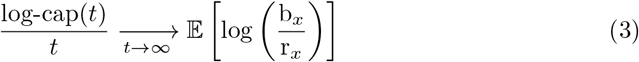

in probability. From this, we define the long-term increase of the log-capital as :

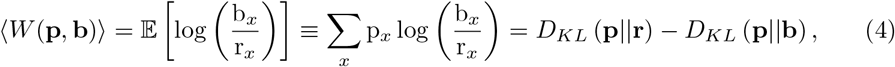

where *D*_KL_ stands for the Kullback-Leibler divergence [2, sec. 2.3]. From an information theoretic point of view, (3) and (4) imply that the capital of the gambler increases in the long term only if the gambler has a better knowledge of **p** than the bookmaker, otherwise it decreases.

It also follows from this analysis that the optimal strategy **b**^KELLY^ = **p**, called *Kelly’s strategy* [1], overtakes any other strategies in the long-term. Its optimum growth rate is the positive quantity *D*_KL_ (**p‖r**). When the odds are fair, the strategy **b**^NULL^ = **r** also plays an important role. We have called this strategy the *null strategy* [16], because it yields asymptotically a constant capital as can be seen from Eqs. (3)-(4).

Risk can be estimated using the volatility, i.e. the asymptotic variance of the fluctuations of the growth rate of the capital. It is well known that this measure is not perfect and it is less appropriate compared to methods that account for the asymmetry in the direction of fluctuations, since positive fluctuations of gain with respect to the mean, are not risk, while negative fluctuations are. In section 4, we will explore an alternative measure of risk precisely for this reason but for the moment, let us use the volatility which allows for tractable calculations.

Since we have considered independent races, by the central limit theorem, the rescaled log-capital converges in law towards a centered Gaussian distribution of unit variance :

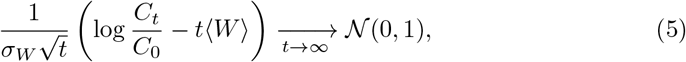

Where

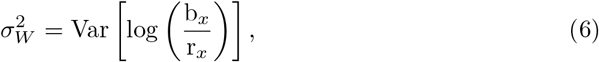

is the volatility. From this definition, one can see that the *null strategy* has a zero volatility, *i*.*e*. it is risk free. Note also that for an arbitrary strategy, risk is relevant at intermediate times, *t* « (*σ*_*W*_ */ ⟨W*⟩)^2^, long-enough for the central limit theorem to apply but not too long for deviations from exponential growth to become negligible.

## 2 Game-theoretic formulation of the asymptotic growth rate

Here, we recall some key results of a game-theoretic formulation of the asymptotic growth rate [22], first in Kelly’s case (diagonal odds) and then in the case where the odds are not diagonal. From the point of view of game theory, Kelly’s model is a zero-sum multiplicative game [23, 24]. Indeed, the average growth rate is the pay-off; the first player is the gambler, because he/she attempts to place the bets **b** so as to optimize the growth rate, while the second player controls horses probabilities **p**. In biological applications, the second player could represent a fluctuating environment. One could object that in real life, the environment is not trying to minimize the growth rate of the system as the second player in the game setting would, nevertheless this case is meaningful since it corresponds to a worst case for the gambler as explained below.

### 2.1 Kelly’s case (diagonal odds)

For Kelly’s optimal strategy **b**^KELLY^ = **p**, the growth rate is

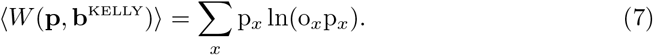

Let us now evaluate the worst possible scenario with the given odds, i.e. the value of **p** such that his/her growth ra is minimal. This can be done by minimizing the function Ψ(**p**) = *(W* (**p, b**^KELLY^)*) − λ* Σp_*x*_ with respect to **p**, where the Lagrange multiplier enforces the normalization of **p**. One obtains that the worst scenario occurs when

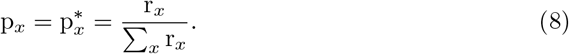

One can then write

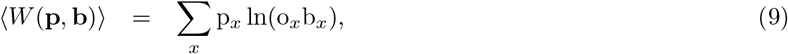

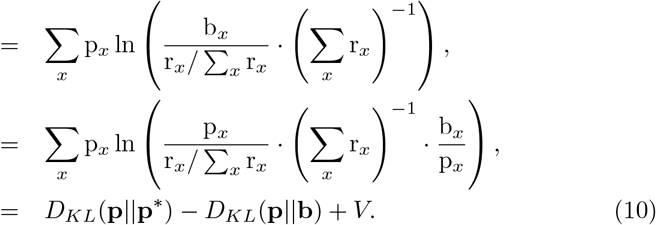

These three terms have the following interpretations:

- *D*_*KL*_(**p‖ p**^*∗*^) is the “pessimistic surprise”, due to the realization that things are not as bad as they could be.
- −*D*_*KL*_(**p b**) is the “better’s regret”, due to the realization that the gambler plays sub-optimally.
- *V* = − ln Σ_*x*_ r_*x*_ is the value of the game, the minimal growth rate that the gambler can expect irrespective of how **p** is chosen. In practice, this minimum is attained for the *null strategy* where **b** = **p**^*∗*^. Further, when the odds are *unfair, V <* 0, whereas when the odds are *super-fair, V >* 0. This definition of the *null strategy* generalizes the previous one when the odds are not fair.

### 2.2 Non-diagonal odds

In the general case, the matrix of the odds o_*xy*_ that gives the reward to a bet *y* when the winning horse is *x*, is non-diagonal, and the corresponding growth rate may be written as :

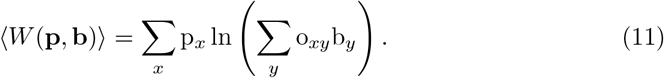

A particular case of non-diagonal odds corresponds to the situation described in the original Kelly’s paper as ‘track take’, where the gambler has the option to not bet a fraction of his/her capital. In that case the growth rate is

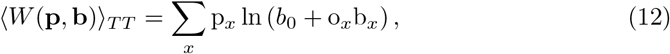

where *b*_0_ is the fraction of the capital which is not bet. This case corresponds to a non-diagonal matrix of odds which contains a diagonal part and an isolated full column filled with ones. The optimal solution for the bets has been considered in the original Kelly’s paper. This solution can be recovered with the Karush-Kuhn-Tücker (KKT) method as shown explicitly in Ref. [25].

This extension of Kelly’s model to non-diagonal odds is particularly important for applications to biology because a given phenotype is never only adapted to one one specific state of the environment, instead there is a distribution of environment states that corresponds to a given phenotype (the equivalent of the bets).

An explicit solution of the optimization with respect to the bets can be obtained provided two conditions (i) and (ii) are met for the odds matrix. The first condition (i) requires that this matrix is invertible and simplex preserving. Simplex preserving implies that multiplying any probability vector by the inverse matrix will keep the vector inside the simplex. This means that the odds matrix, viewed as a game, is *fully mixing* [23]. An equivalent mathematical condition is that the inverse-odds matrix r = o^*−*1^ has no zero in it. When this condition is satisfied, we can build the following game-theoretic solution:

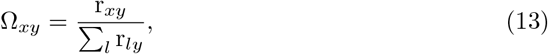

which is such that

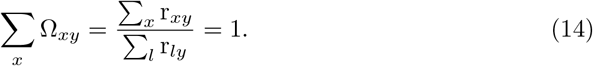

Then, one can show that the optimal bets are given by

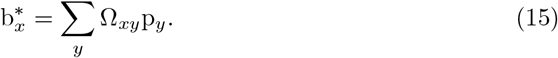

In order to proceed with the game-theoretic analysis, we need to look for the worst case scenario, i.e. for the value **p**^*∗*^ of **p** yielding the minimal growth rate for the optimal strategy of the gambler. Using again the method of Lagrange multiplier, one finds

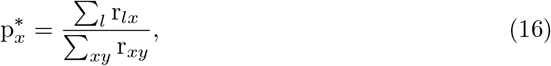

which is acceptable provided all the components of *p*_*x*_ are non-negative. This requires that for all *x*, Σ_*l*_ r_*lx*_ *>* 0, which is our second condition (ii). When both conditions hold, the matrix Ω is stochastic (or more precisely pseudo-stochastic because it can contain negative elements), and there is unique pair (*p*^*∗*^, *b*^*∗*^), which represents a Nash equilibrium for the matrix game defined by the odds matrix *o*.

To obtain the equivalent of the decomposition of Eq. (9) for the case of non-diagonal odds, we start by evaluating the optimal growth rate when the bets are optimal and given by Eq. (15):

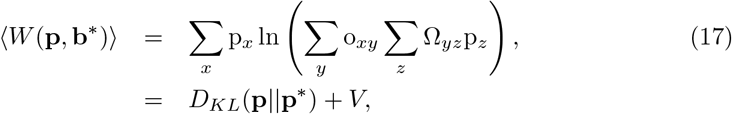

where **p**^*∗*^ is the one of Eq. (16) and now the value of the game is *V* = ln(Σ_*xy*_ r_*xy*_). In a second step, one can then check that the growth rate for non-optimal bet **b** is less or equal than *W* (**p, b**^*∗*^) and that the difference is the term associated to the better’s regret. In the end, one finds that the decomposition of the asymptotic growth rate of Eq. (9) still holds for non-diagonal odds, provided one takes into account the new definitions of of **p**^*∗*^, ⟨*W*⟩ and *V*, and conditions (i)+(ii) hold. In particular, the value of the game still has the same interpretation and the minimum of the growth rate is still attained for the *null strategy*.

### 2.3 Illustration with a three horses example

Let us illustrate this game-theoretic framework for a simple case of three horses in which only the gambler can play optimally. Let’s consider specifically a diagonal and a non-diagonal odds matrix given by:

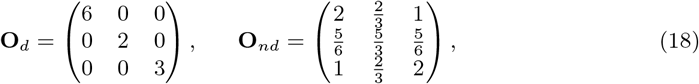

and an environment characterized by the vector *p* = (*p*_0_, *p*_1_, *p*_2_) where *p*_0_ is varied in (0, 1), 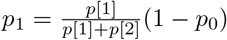 and 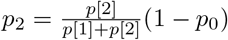, and (*p*[1], *p*[2]) = (0.5, 0.3).

When the game is non-fully mixing, there is an analytical solution for the optimal solution namely (13)-(16). When it is not, we need to resort to numerical optimization. Simulated annealing and KKT optimization are two possible methods to do this. We have found empirically that the later gives better results than the former for low dimensions problems, which is the case here, since we only consider three horses. Below, we only use the KKT method. The optimization problem we want to solve is :

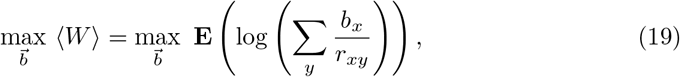

subject to

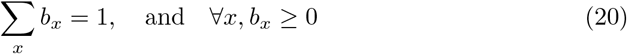

To solve it with KKT method, one introduces the functional :

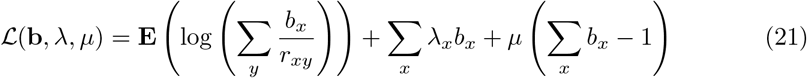

where *λ*_*x*_ and *μ* are Lagrange multipliers. Since the problem is concave for **b**, we set the first derivative to zero to obtain the point of maximum, i.e. the optimal strategy **b**^*∗*^.

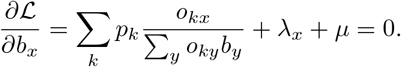

The KKT solution is built by combining the solution of that equation for three mutually exclusive cases : (i) one bet *b*_*i*_ is zero while the other two *b*_*j*_ for *j* ≠ *i* are strictly positive. In that case, we have *λ*_*i*_ *>* 0. (ii) two bets are zero *b*_*i*_ = *b*_*j*_ = 0 while the last one is strictly positive. In that case, similarly *λ*_*i*_ *>* 0 and *λ*_*j*_ *>* 0, and (iii) none of the bets is zero, which is the same solution as that of (13)-(16).

A comparison between the result of KKT optimization (symbols) with the game-theoretical solution of equation (13)-(16) is shown in Fig. (1) which shows the optimal strategies 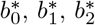 as a function of *p*_0_ ∈ (0, 1), for the case of diagonal odds (a) and non-diagonal odds (b). It is interesting to notice that, as the environment probability vector varies, only in a certain interval of values of *p*_0_ the game is non fully-mixing and (13)-(16) apply. Outside of this interval, one or more *b*_*i*_ = 0, the game is non fully-mixing and the game-theoretical solution is no longer correct. This explains the deviation with the correct result obtained from the KKT maximization for small value of *p*_0_.

**Fig 1.**
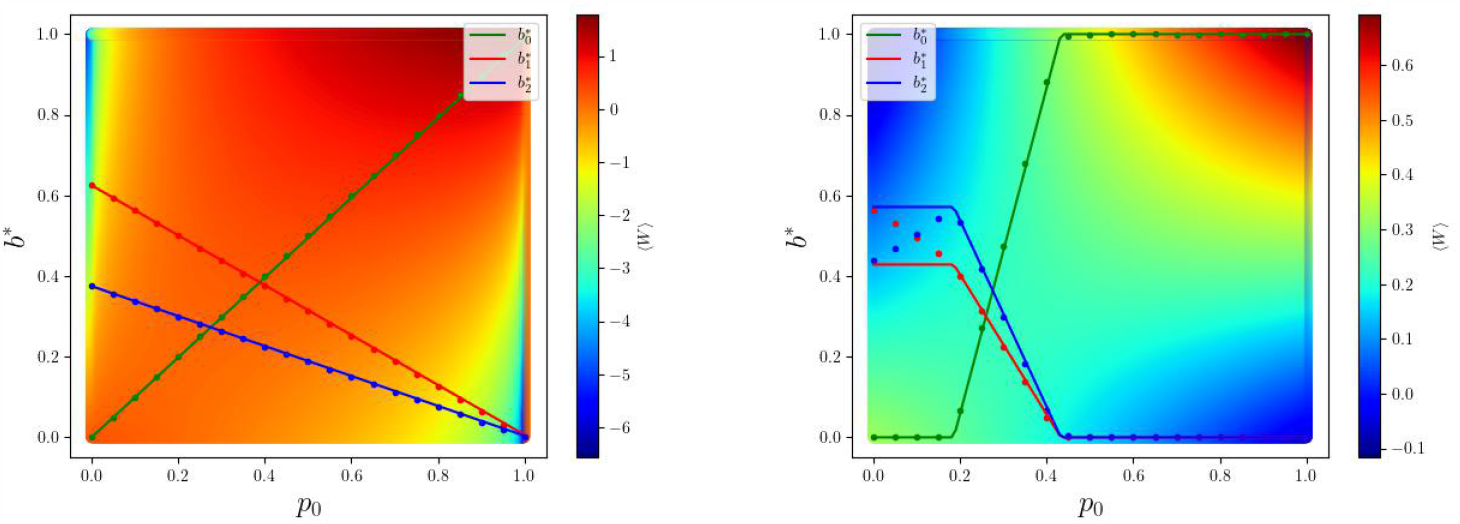
KKT solution vs. game theoretic solution. Optimal bets 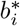 on the three horses from KKT optimization (symbols) and game-theoretical solution deduced from (13)-(16) (solid lines) as function of the probability on the first horse *p*_0_. Figure (a) corresponds to the diagonal odd matrix **O**_*d*_, figure (b) to the non-diagonal odd matrix **O**_*nd*_, the green color corresponds to first horse, red color for the second one, blue for the third one. The colored intensity plot displays the average growth rate.

The figures also show the average growth rate as a color plot, projected along the plane (*p*_0_, *b*_0_) for the diagonal and non-diagonal cases. Note that in this color plot, a particular choice is made for varying the parameters *p*_1_, *p*_2_, *b*_1_, *b*_2_ as *p*_0_ and *b*_0_ vary while satisfying normalization constraints. In the diagonal case, one can see that the growth rate is the highest for a fixed *p*_0_, *i*.*e*. along the green line where *b*_0_ = *p*_0_ as expected from Kelly’s gambling. Moreover, in the non-diagonal case, the game-theoretical solution takes instead the form of piece-wise linear functions. The color plot for the average growth rate shows the presence of a saddle point, which is visible here only in projection.

### 2.4 Illustration of the reduction of a game

The derivation of the game-theoretical solution of equation (13)-(16) assumes (i) a fully mixing game, (ii) an odd matrix which is invertible and simplex preserving. We have seen in the previous section case what happens when the game is non-fully mixing. In that case, the strategies corresponding to zero bets or zero probability *p*_*i*_ become in a sense irrelevant because they are dominated by other strategies corresponding to non-zero *b*_*i*_ or *p*_*i*_. These irrelevant strategies can be removed, and when doing so, one transforms the game into what is called the essential part of the game [23]. Let us illustrate this reduction procedure by starting with the 3×3 game defined by **O**_*nd*_, which we will reduce to a 2×2 game. The reason that we do not consider a reduction to a 3×2 game for instance, is because we need the reduced game to be a square matrix so that it can be invertible and (13)-(16) can apply.

Let us consider an input strategy vector for the environment given by (*p*[1], *p*[2]) = (0.8, 0), so that while *p*_0_ varies, *p*_2_ = 0 always, and one can check that in that case one also has 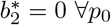. As a result, the last horse should never be played and the odd matrix can be reduced by removing the last row and last column in **O**_*nd*_. The odd matrix of the essential part of the game is then:

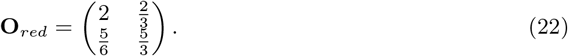

Now, since the essential part of the game is fully-mixing, and fulfills the assumptions under which relations (13)-(16) have been derived, one can use these equations to obtain the optimal solution and check that they agree with KKT solution as shown in Fig. 2.

**Fig 2.**
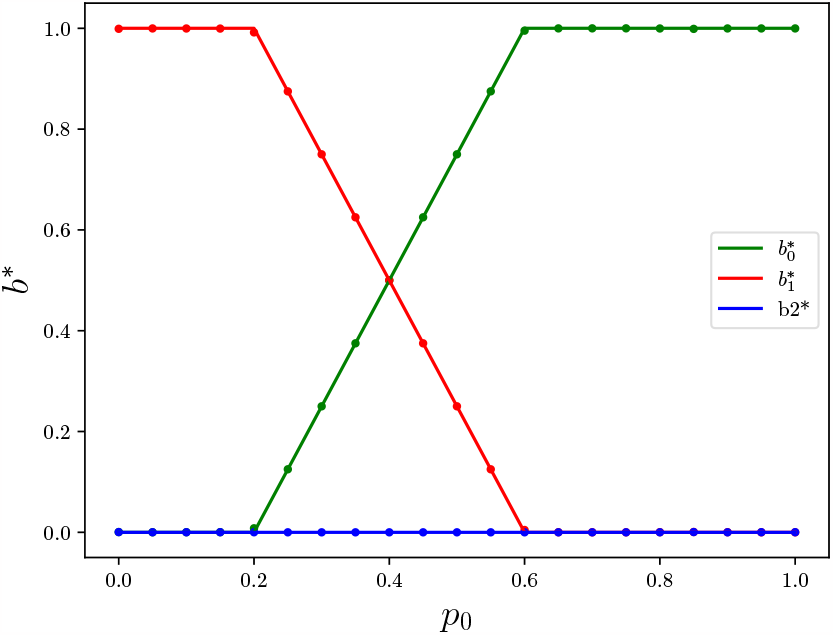
Illustration of the reduction of the game to its essential part. Comparison between the result of KKT optimization (symbols) and the game-theoretical solution (solid lines) for the reduced odds matrix **O**_*red*_, which is the essential part of the game defined by **O**_*nd*_.

## 3 Risk-return inequalities and their associated trade-off

### 3.1 Non-fair but diagonal odds

In recent work, we have studied the trade-off between the mean growth rate and the risk, measured by the volatility in the case of Kelly’s original model with *fair* (and diagonal) odds [16]. This trade-off is embodied in the following mean-variance inequality :

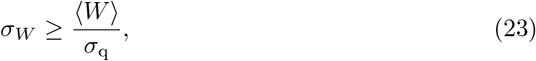

where *σ*_*W*_ is the volatility, ⟨*W*⟩ the average growth rate and *σ*_*q*_ is the standard deviation of a distribution q_*x*_ = r_*x*_*/*p_*x*_ which compares the probability of races outcomes with the risk-free strategy. Inequality (23) applies for any *(W) >* 0. In the negative growth region, it only applies near the null strategy where *D*(*r*‖*b*) ≈ 0. This inequality means in practice that a capital growing exponentially with a rate ⟨*W*⟩ > 0 necessarily has a non-zero risk measured by the volatility. Recently, a similar bound has been derived for a wide class of financial models such as the Black-Scholes and the Heston models [26] by building on previous work on the so called thermodynamic uncertainty relations of Stochastic Thermodynamics.

Let us first generalize this result to the case where the odds are not *fair* but still diagonal. To do that, we introduce a new distribution similar to that of the inverse-odds, but which is normalized. An obvious choice is the distribution 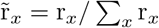. Further, the strategy 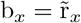 corresponds to the *null strategy* introduced previously. For such a strategy, the asymptotic growth rate *W* is equal to *V*, independently of the choice of the bets and of the horse winning probabilities.

The definition of the distribution q is unchanged with respect to the case of fair odds, the only difference is that now ⟨q⟩ ≠ 1. Let us now go through the same steps which lead previously to Eq.(23). We start with

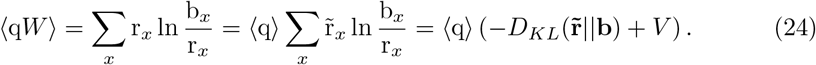

Now, we write the covariance between q and *W* as :

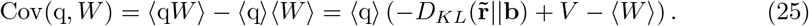

Using Cauchy-Schwartz inequality, namely 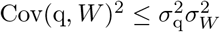 and the positivity of the Kullback-Leibler divergence, we obtain the generalization of Eq. (23) for non-fair odds as :

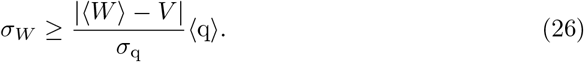

valid for any ⟨*W*⟩ − *V >* 0 and only close to the null strategy for ⟨*W*⟩ − *V <* 0.

From the inequality of Eq. (26), we find that any strategy different from the *null strategy* which is risk free will have a non-zero volatility *σ*_*W*_. Note also that the inequality is saturated for the risk free strategy for which *σ*_*W*_ = 0 and ⟨*W*⟩ = *V*. A numerical illustration of that inequality is provided in figure 3 for a case where *V* = 0 and a case where *V >* 0.

**Fig 3.**
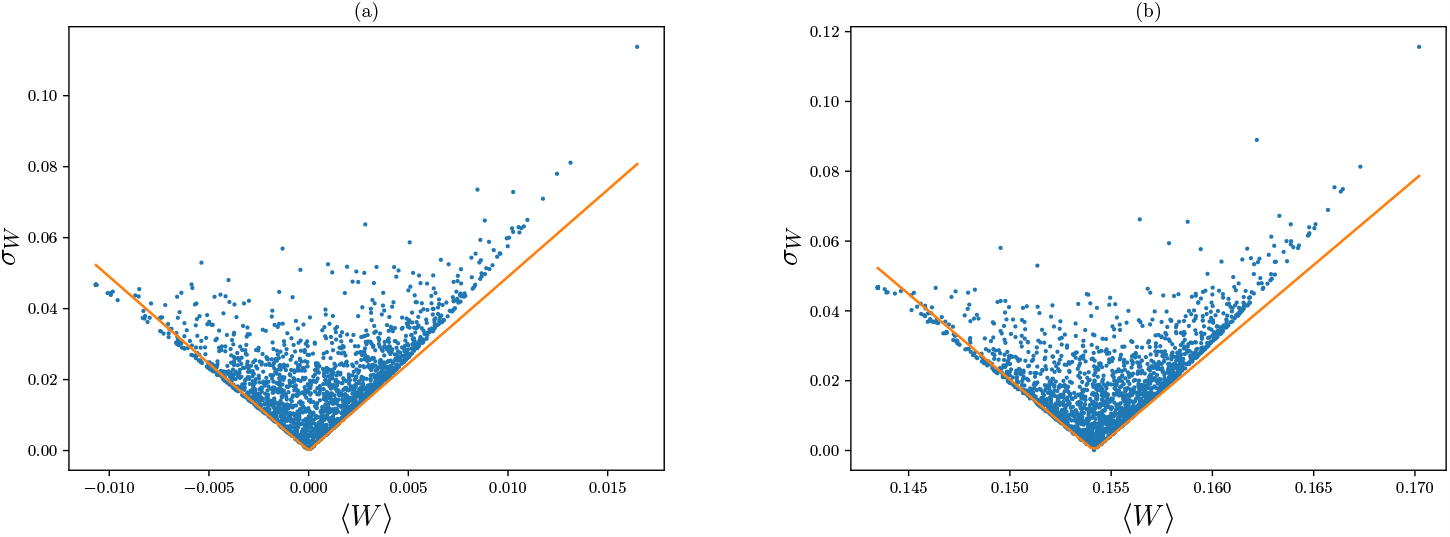
Pareto plots for fair and super-fair odds in the variables volatility *σ*_*W*_ versus growth rate *W*. The cloud of points displays an ensemble of feasible random strategies for non-diagonal (a) fair and (b) super-fair odds, where *V >* 0. The solid lines test the inequality of Eq. (26), which is globally valid to the right of the risk-free strategy (where both lines meet) but just locally valid to the left of the risk-free strategy.

### 3.2 Non-diagonal odds

The mean-variance trade-off for non-diagonal can be derived similarly to the diagonal case, provided the same conditions denoted (i) and (ii) hold. Condition (i) is the invertibility of the odds matrix, assuming it holds, we call *r* the inverse-odds matrix.

Now, the relevant q distribution has the form q_*x*_ =Σ _*y*_ r_*yx*_*/*p_*x*_ so that ⟨q⟩ = Σ_*xy*_ r_*xy*_ = exp(*−V*) in terms of the value of the game *V* = *−* ln(Σ_*xy*_ r_*xy*_).

We start again with

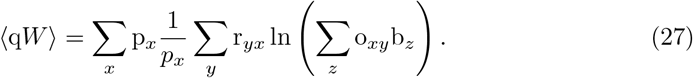

In order to write this term as a KL divergence, we introduce two new normalized distributions :

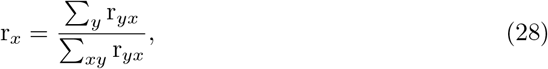

which is acceptable as a distribution provided condition (ii) holds. Similarly, we introduce

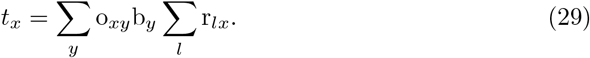

It is easy to see then that

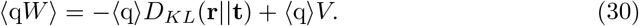

In the end, we obtain the same relation as in Eq. (26), provided one takes into account the new definitions of the distribution q, ⟨*W*⟩ and *V*, and conditions (i)+(ii) hold.

### 3.3 Further consequences of the risk-return trade-off

When the odds are fixed, the clouds of points of Figs. 3 change when the probability vector of the horses to win (the **p** vector) changes as shown in Fig. 4. Each choice of this vector generates a separate cloud of points, and all these clouds of points have the same lowest point in common, namely the risk free strategy, where ⟨*W*⟩= *V* and *σ*_*W*_ = 0 independently of the **p** vector. Each cloud of points admits a tangent vector near this risk free strategy with a slope determined by an inequality of the form (26). There is a different slope for each tangent since the slope depends on the **p** vector. Now, if some information is known about the family of distributions of (the **p** vector), one can combine all these bounds to obtain a general bound on all the possible values of the slopes. Such a global bound would then inform on the minimum level of risk irrespective of the distribution **p**.

**Fig 4.**
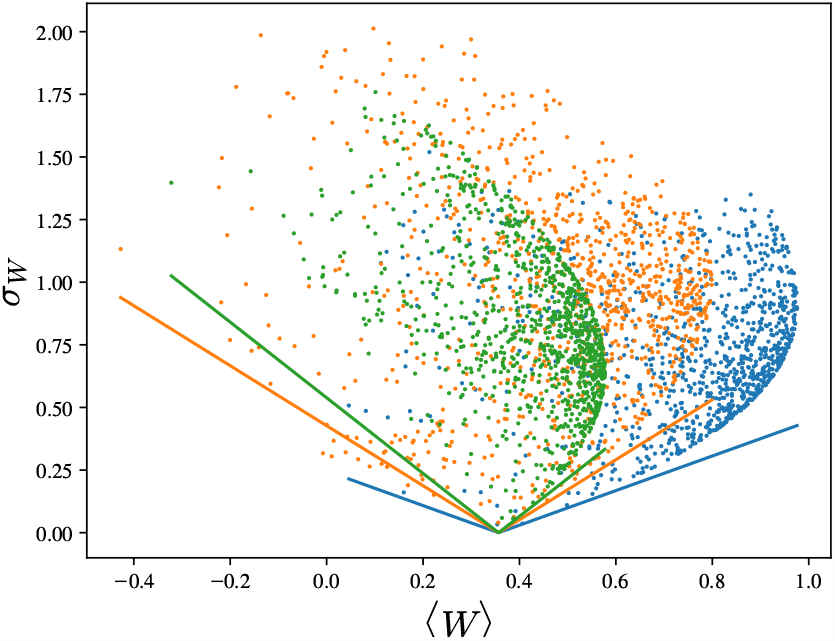
Clouds of points from Kelly’s horse race model in the plane (*σ*_*W*_, ⟨*W*⟩) for various probability p vectors. The value of the game is unchanged as it is independent of the **p** vector, here it is *V* = 0.36. The solid lines have the same meaning as in Fig.3.

## 4 Risk quantification beyond volatility

### 4.1 Extinction probability for geometric brownian motion

Alternative measures of risk beyond volatility are needed because the volatility is symmetric, i.e. it describes positive or negative fluctuations. Therefore, it does not conform to the intuitive notion of risk, which is asymmetric since it is only related to negative fluctuations. To build a more appropriate measure of risk, we turn to a continuous approximation of the trajectory of the log-capital as a geometric Brownian motion. This corresponds to the asymptotic regime for the central limit theorem of Eq. (5), in which the log-capital is distributed according to a normal law with ⟨*W*⟩ as mean and *σ*_*W*_ as standard deviation. Assuming the log-capital *y*(*t*) = log *C*_*t*_ starts from an initial value *y*_0_, the probability that it reaches the value *y* at time *t*, namely 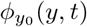 is:

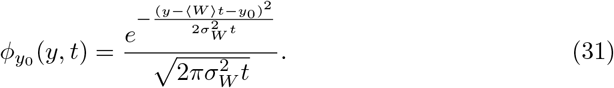

Then, the extinction probability is defined as the probability that the log-capital reaches a certain low threshold at any time *t*^*’*^ *< t* for the first time. Further, 𝒫(*t*) = 1 − 𝒮(*t*), where 𝒮(*t*) denotes a survival probability, defined as the probability that the log-capital *y*(*t*) did not ever reach the low threshold *l at any time t*^*’*^ *< t* assuming that it started with the value *y*_0_ at time 0 with *y*_0_ *> l*.

The survival probability *𝒮*(*t*) can be evaluated from the classic image method. According to this method, one writes the probability *P* (*y, t*) for the random walker to reach *y* at time *t* as a linear combination of 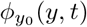 and *ϕ*_*m*_(*y, t*) where *m* is the position of the image. By enforcing the condition *P* (*y* = *l*) = 0 at all times, one finds *m* and an explicit form for *P* (*y, t*). Then, the survival probability is 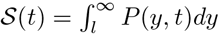. One obtains

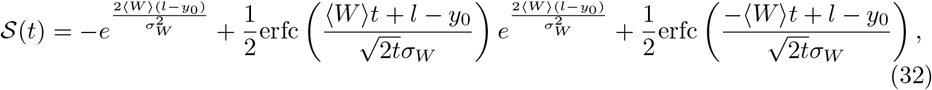

where erfc(x) denotes the complementary error function (i.e. erfc(x) = 1 − erf(x), where erf(x) is the error function). It is straightforward to check that *𝒮*(0) = 1. One also finds that 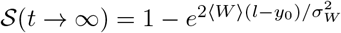. Therefore, it is a meaningful survival probability provided ⟨*W*⟩*>* 0 if *l < y*_0_ or ⟨*W*⟩ *<* 0 if *l > y*_0_. Let us focus on the case ⟨*W*⟩ *>* 0, for which the capital is growing exponentially on long times. The larger ⟨*W*⟩ *>* 0 or the higher the distance between the starting point *y*_0_ and the threshold *l*, the less likely the log-capital reaches the low threshold, as one would expect. Since a negative fluctuation of the capital is needed to reach this low threshold, such an event can only occur at rather short times because at long times the capital is growing exponentially as illustrated in Fig 5a. Further, it can be shown that the event is guaranteed to occur when 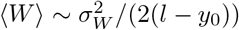.

**Fig 5.**
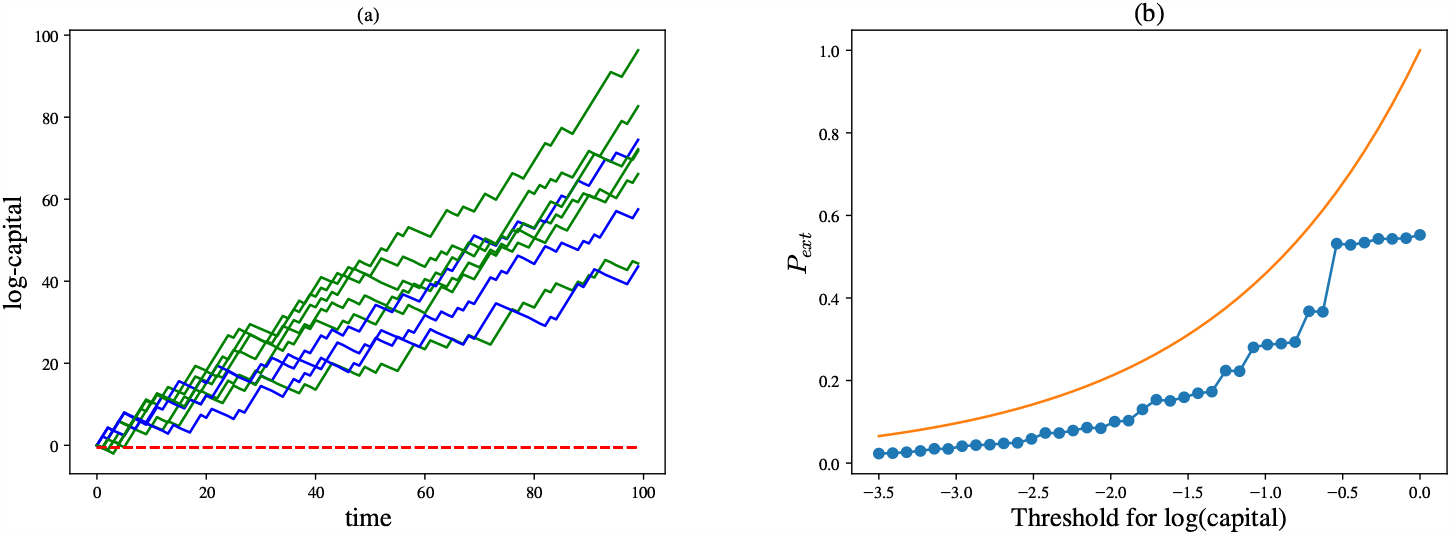
(a) Trajectories of Kelly’s horse race model and (b) comparison between the extinction probability in Kelly’s model and in the geometric Brownian motion that matches the parameters of Kelly’s model. For figure (a), trajectories either never reach the target (blue solid curves) or do reach it (green solid curves), typically at short times. The threshold is set at *l* = − 0.5 (red dashed line). For figure (b), the extinction probability is computed for Kelly’s model vs. position of the threshold after 100 races, and in the long time limit for geometric Brownian motion. In both figures, horse probabilities and returns are *p* = (0.36, 0.15, 0.49) and *r* = (0.63, 0.31, 0.06).

From these considerations, an inequality similar to that of Eq. (23) can be derived to describe the mean growth rate-risk trade-off using the extinction probability

*𝒫*_*ext*_ = *𝒫*(*t → ∞*) as a proxy of risk instead of the volatility. From the expression of *𝒮*(*t → ∞*) above, it is straightforward to obtain in the case of fair odds and when ⟨*W*⟩ *>* 0:

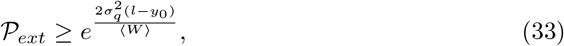

which shows that in order to reduce risk (as measured by extinction probability), one needs to reduce the growth rate or bring the threshold further away from the initial capital as one would expect.

Further characterizations of risk could be considered. For instance, the distribution of first passage times for the log-capital to reach the threshold can be obtained from the opposite of the time derivative of *𝒮*; and using more advanced arguments, one can also compute analytically the distribution of the time where the log-capital reaches its maximum for an arbitrary value of the drift. This question has been studied in finance because it is related to the optimization of the time to sell/buy a stock [27].

In Fig. 5b, we compare the extinction probability *𝒫* (*t*) for a fixed final time *t* as function of the threshold value *l*, for Kelly’s horse race and for its approximation using geometric Brownian motion. In the case of Kelly’s model, many stochastic trajectories are simulated from the model in the same conditions and from the statistics of these trajectories an empirical estimation of the extinction probability *𝒫*(*t*) is obtained. The simulation results of Kelly’s model displays steps, which follow the trend given by the continuous model. The presence of these steps can be traced back to the fact that in Kelly’s model the log-capital changes by discrete increments at discrete time intervals. In Fig. 5b, one sees a comparison between the extinction probability evaluated from Eq. (32) using geometric brownian motion with a simulation of that quantity evaluated using Kelly’s model. As shown in the figure, the prediction of geometric brownian motion is very close to that of Kelly’s model in the left part of the figure where the threshold takes its minimum value. This is expected because in this regime the trajectory contains a large number of steps to reach the threshold and therefore the continuum approximation is well verified. In contrast, this does not happen on the right part of the figure, where the discreteness of Kelly’s model is quite apparent.

### 4.2 Risk-constrained Kelly gambling

In our first study of risk-constrained Kelly gambling [16], we have introduced a penalization proportional to the volatility in the optimization of Kelly’s growth rate with respect to the bet vector. This was done with the following objective function, which interpolates between the maximization of the growth rate and the minimization of the variance of the growth rate :

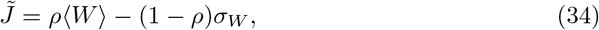

with 0 ≤ *ρ* ≤ 1. In this approach, the parameter *ρ* plays the role of a risk aversion parameter, and the optimal bets are parametrized by it. From this optimal bets, one can build Pareto diagrams that represent the minimum amount of fluctuations for a given growth rate. An example of these Pareto diagrams is shown in Fig. 6a.

**Fig 6.**
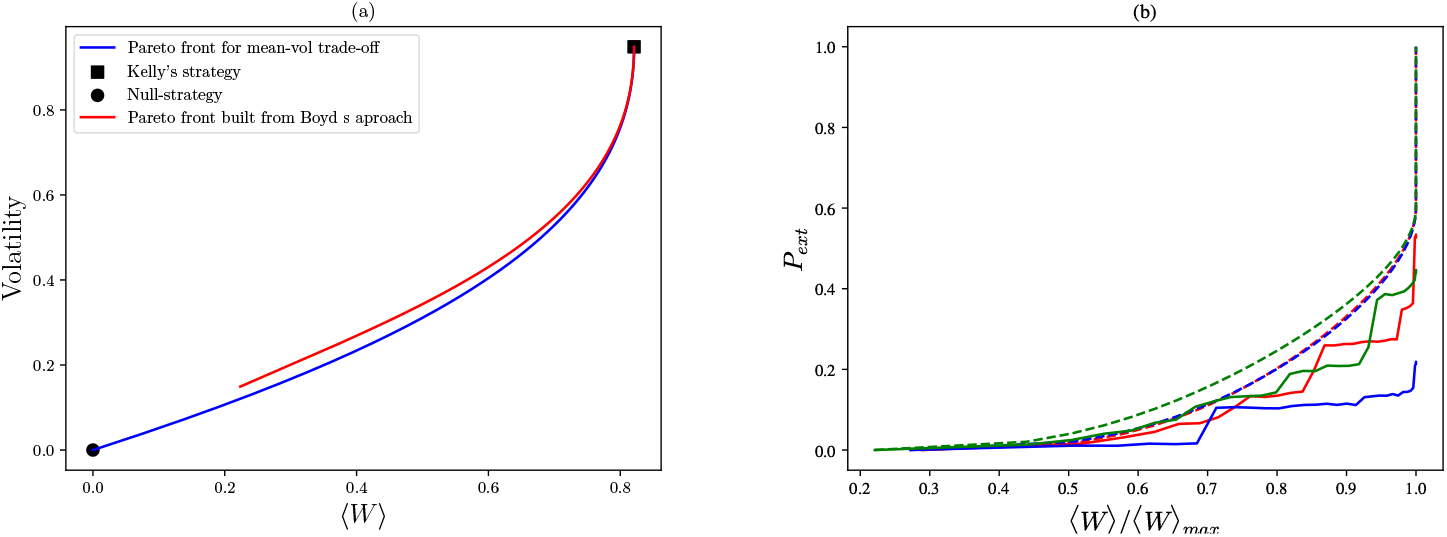
(a) Pareto front of the volatility versus the average growth rate, and (b) extinction probability as a function of (normalized) growth rate for Boyd’s optimization (solid lines). In figure (a), curves are calculated according to the mean-volatility trade-off approach (solid blue line) and volatility vs growth rate line for Boyd’s optimization of growth rate with extinction probability constraint (solid red line). In figure (b), the three different sets of parameters are: blue line is computed with the same horse probabilities and returns as in (a), red and green show two other different combinations taken at random. Dashed lines correspond to Boyd’s method bound *β* for extinction probability (*P* (*C*_*min*_ *< α*) *< β*). Each extinction probability curve is bound by the dashed line of matching color. Extinction probability is computed from 40000 simulations of 100 races for each value of the growth rate shown. For both plots *α* = *l* = 0.6

Instead of using the volatility to constrain the growth rate in Kelly’s gambling, another approach is to introduce a constraint into the optimization of the growth rate to enforce that the extinction probability does not go beyond a certain threshold [28]. As usual, the constraint is taken into account with a Lagrange multiplier. To properly define that approach, it is convenient to introduce :

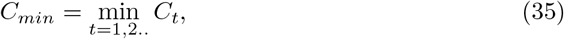

which represents the lowest value reached by the capital *C*_*t*_ during the observed time before it goes on increasing. The drawdown risk is quantified by the probability that this minimum goes below a target value, *P* (*C*_*min*_ *< α*), where *α* is the target value for the capital. Then the constraint on the probability of drawdown has the form *P* (*C*_*min*_ *< α*) *< β*. For example, we might take *α* = 0.7 and *β* = 0.1, meaning that we require the probability of a drawdown of more than 30% to be less than 10%. This drawdown risk does not have in general a simple form as function of the bet vector, it can only be obtained numerically by solving a non-linear optimization problem with non-linear constraints. While this optimization problem is difficult, Boyd et al. introduced a bound on the drawdown risk that results in a tractable convex constraint [28]. This bound reads as follows:

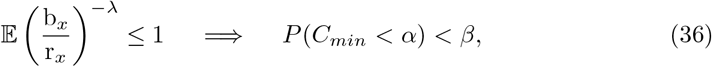

where *λ* is defined as

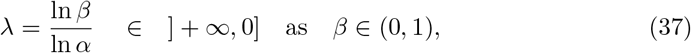

which means that, by varying the maximum extinction probability allowed, hence varying *λ*, our optimization is more or less sensitive to risk. In that sense, *λ* is a risk aversion parameter similar to *ρ* in the mean-variance approach.

In the following, we fix the value of *α*. We consider the case of three horses, with an initial capital *C*_0_ = 1, and we use *p* = (0.1, 0.2, 0.7) and *r* = (0.7, 0.1, 0.2). With these values, we obtain the optimal strategy **b**^*∗*^ for different *β ∈* (0, 1). Once the optimal strategy is obtained for a fixed *β*, we can compute the growth rate ⟨*W*⟩ and the variance *σ*_*W*_ for that particular strategy. Hence, we obtain the diagram in the coordinates (⟨*W*⟩ − *σ*_*W*_) shown in Fig. 6a.

In this figure, we observe that the two measures of risk lead to comparable plots. Further, the blue line is always below the red line which is expected since the blue plot represents the set of points where variance is minimized for a given growth rate. At Kelly’s point, both curves meet since this corresponds to the case *β* = 1 for which the Boyd’s approach reduces to the simple optimization of the growth rate as done in Kelly’s approach. We have observed that these features are robust with respect to the choice of *α*. Note that the red curve from Boyd’s approach does not reach arbitrary low values of the growth rate because of the choice of the lowest value of *β*. In Boyd’s approach, the null strategy is only reached asymptotically as *β* approaches zero.

In Fig. 6b we analyze the bound *β* on the actual extinction probability. Using simulations, we computed the probability of extinction for the optimal bets *b*^*∗*^ obtained by Boyd’s maximization. These simulations where run for the same parameters considered in Fig. 6a and for two other sets of horse probabilities and returns chosen at random. Finally, we show both the probability of extinction and the corresponding bound *β* (in matching color) as a function of average growth rate. As apparent from the figure, the probability of extinction is always below its bound, but depending on the parameters chosen it may be tighter or looser. Although these curves do not represent a Pareto front for probability of extinction and growth rate, they show that in general probability of extinction increases with growth rate, making Kelly’s the riskiest strategy.

We observe sudden increases in the extinction probability as a function of the growth rate at some points. This may be related again to the discreteness of the log-capital as in figure 5, where we see steps in probability of extinction when the threshold is modified. In fact, through the optimization procedure, the average gain and the threshold are connected, so that one can parametrize the optimal solution with the threshold or the gain as in figure 6b.

It is interesting to notice that above a certain value of *β* = *β*^*∗*^, the curve for the bound becomes vertical in Fig. 6b: Kelly’ s strategy is always the optimal strategy when the bound imposed on the probability of extinction becomes high enough. This behavior is akin to a phase transition separating an optimal solution which is Kelly’s like from a non-Kelly strategy. Indeed, when the constraint 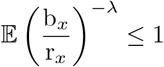 is inactive for specific values of *p* and *r*, the solution of the optimization is the one without the constraint, i.e. Kelly’s solution.

It is easy to check that in the region 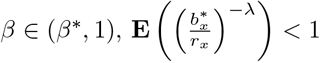, for the values of *p* and *r* chosen above. There, Kelly’s strategy is always optimal in this interval of *β* values where the probability of extinction is high. Note that the lower end of the vertical line corresponds to the case where the constraint becomes active and Kelly’s strategy no longer fulfills the condition, so the optimal strategy then becomes different to Kelly’s betting, in order to lower the risk.

In some specific cases, the vertical line does not exists (*r*_*i*_ *> p*_*i*_ ∀*i*) which corresponds to unfair odds, or the plot shows only the vertical line corresponding to Kelly’s regime for all *β* when *r*_*i*_ *< p*_*i*_ ∀*i*.

In the case where the vertical line does not exists, odds are unfair for the gambler, who should avoid Kelly’s strategy because it leads to a high extinction probability (which means high probability of bankruptcy for the gambler). In other words, when conditions are not favorable (in terms of the odds or of the distribution of the probabilities of the environment, gamblers (respectively biological systems) can not maintain themselves at Kelly’s point except at the cost of a large extinction probability of the population (respectively bankrupcy probability). Instead, in good conditions for growth, Kelly’s strategy is optimal.

## Conclusion

In this article, we have explored an extension of Kelly’s model to the case non-diagonal odds, an extension which is particularly relevant for biological applications. We have analyzed in this more general setting, the trade-off between the average growth rate and the volatility, which is known in the financial literature as risk-return trade-off. We have also explored an alternate measure of risk beyond volatility, namely the extinction probability, which can easily be calculated if the races are uncorrelated as a realization of geometric brownian motion. Our main result is that this measure of risk leads to comparable results as obtained with the volatility as far as the risk-return trade-off is concerned. In particular, the inequality that embodies the trade-off between average growth rate and volatility can be expressed similarly as an inequality in which the extinction probability replaces the volatility. Given the derivation of these inequalities, which mainly follow from information-theory, we expect that they should hold in a broader context whenever growth can be characterized with a multiplicative process.

One area of applications of these ideas concerns the adaptation of microbial populations to uncertain environments. In this context, E. van Nimwegen et al. recently introduced the notion of growth rate dependent stability (GRDS) which allows cells to stabilize their phenotypes at high growth rate [29]. As a result, cells implementing GRDS have a fitness advantage with respect to those using only the standard bet-hedging strategies. We note that such a stabilization of the phenotypes by GRDS reduces the fluctuations of the growth rate, which is a way to overcome the negative impacts due to the trade-off between growth rate and risk studied in this paper.

Another area of application of these ideas is ecology, where one central question is to understand what determines the diversity of species and the coexistence between species. In this context, biodiversity can be regarded as a form of biological insurance against disruptive effects of the environment, because biodiversity reduces the variability in ecosystem properties that arises due to differential responses of species to environmental variations [30]. This line of research supports the idea that there is in ecology a trade-off similar to the one between growth and risk. The question of the persistence of species is also related to that trade-off. Indeed, persistence of species can not be decided based only on the growth rate as in model coexistence theory, fluctuations matter too, where fluctuations can either refer to that of species abundances [31] or that of the growth rate itself [32]. In the later case, this agrees with the choice of variables used in this paper. In fact, we see that both in Ref. [32] and in our work the crucial quantity is the ratio between the growth rate and the standard deviation of the fluctuations, which is precisely the ratio that enters in the inequality we have derived and which is known in finance under the name of Sharpe ratio.

In this paper, we also explored how ideas of game theory can be used together with Kelly’s model by building on Ref. [22]. We found that when the game is not fully mixing for certain environmental probabilities, it can be reduced to a smaller game, known as the essential part of the game [23], which is fully mixing. This method is interesting because it allows us to break the complexity of the initial problem into the study of a problem of reduced complexity without affecting the optimal strategies.

We hope this work could be useful in analyzing several problems in biology, ecology, econophysics, and social sciences where ideas of bet-hedging and game theory are relevant. For example, in the stock market, the odds matrix that codes for the daily returns from a list of stocks is non-diagonal. However, the challenge is how to deal with the day-to-day randomness in the daily returns themselves.

## Acknowledgments

DL acknowledges support from (ANR-11-LABX-0038, ANR-10-IDEX-0001-02). LD acknowledges support from Spanish Ministerio de Ciencia e Innovación through Grant PID2020-113455GB-I00 and Universidad Complutense de Madrid through “Convocatoria plurianual para la recualificación del Sistema Universitario Español para 2021-2023 (MV24/21)” funded by NextGenerationEU. RP is supported by the Israeli Science Foundation (Grant 776/19).

We acknowledge fruitful discussions with Luca Peliti and Haim Permuter.

## Notes

### Competing Interest Statement

The authors have declared no competing interest.

## References

1. Kelly JL. A new interpretation of information rate. IRE Transactions on Information Theory. 1956;2(3):185–189.

2. Cover TM, Thomas JA. Elements of Information Theory. Wiley Interscience; 2005.

3. MacLean LC, Thorp EO, Ziemba WT. The Kelly Capital Growth Investment Criterion: Theory and Practice. vol. 3. Word Scientific; 2011.

4. Luenberger DG. Investment Science. Oxford Universily Press. Inc; 1998.

5. Proskurnikov AV, Barmish BR. On the Benefit of Nonlinear Control for Robust Logarithmic Growth: Coin Flipping Games as a Demonstration Case. IEEE Control Systems Letters. 2023;.

6. Donaldson-Matasci MC, Bergstrom CT, Lachmann M. The fitness value of information. Oikos. 2010;119(2):219–230.

7. Kussell E, Leibler S. Ecology: Phenotypic diversity, population growth, and information in fluctuating environments. Science. 2005;309(5743):2075–2078.

8. Thattai M, van Oudenaarden A. Stochastic Gene Expression in Fluctuating Environments. January 2004;.

9. Levien E, Min J, Kondev J, Amir A. Non-genetic variability: survival strategy or nuisance? Rep Prog Phys. 2020;84(11):116601.

10. Lacoste D, Rivoire O, Tourigny DS. Cell behavior in the face of uncertainty; 2023.

11. Balaban NQ, Merrin J, Chait R, Kowalik L, Leibler S. Bacterial Persistence as a Phenotypic Switch. Science. 2004;305(5690):1622–1625.

12. Maslov S, Sneppen K. Well-temperate phage: optimal bet-hedging against local environmental collapses. Sci Rep. 2015;5:10523.

13. Cohen D. Optimizing reproduction in a randomly varying environment. Journal of Theoretical Biology. 1966;12(1):119–129.

14. Venable DL. Bet hedging in a guild of desert annuals. Ecology. 2007;88(5):1086–1090.

15. Măgălie A, Schwartz DA, Lennon JT, Weitz JS. Optimal dormancy strategies in fluctuating environments given delays in phenotypic switching. Journal of Theoretical Biology. 2023;561:111413.

16. Dinis L, Unterberger J, Lacoste D. Phase transitions in optimal betting strategies. EPL (Europhysics Letters). 2020;131(6):1–23.

17. Dinis L, Unterberger J, Lacoste D. Pareto-optimal trade-off for phenotypic switching of populations in a stochastic environment. Journal of Statistical Mechanics: Theory and Experiment. 2022;2022(5):053503.

18. Hufton PG, Lin YT, Galla T, McKane AJ. Intrinsic noise in systems with switching environments. Phys Rev E. 2016;93(5):052119.

19. A Despons LP, Lacoste D. Adaptive strategies in Kelly’s horse races model. Journal of Statistical Mechanics: Theory and Experiment. 2022;2022:093405.

20. Rivoire O, Leibler S. The Value of Information for Populations in Varying Environments. Journal of Statistical Physics. 2011;142(6):1124–1166.

21. Tal O, Tran TD. Adaptive Bet-Hedging Revisited: Considerations of Risk and Time Horizon. Bull Math Biol. 2020;82(4):50.

22. Pugatch R, Barkai N, Tlusty T. Asymptotic Cellular Growth Rate as the Effective Information Utilization Rate; 2013. Available from: http://arxiv.org/abs/1308.0623.

23. Barron EN. Game Theory, an introduction. 2nd ed. Wiley. Loyola University, Chicago; January 2010.

24. Giraud G. La théorie des jeux. 3rd ed. Flammarion, Paris; 2009.

25. Smoczynski P, Tomkins D. An explicit solution to the problem of optimizing the allocations of a bettor’s wealth when wagering on horse races. The Mathematical Scientist. 2010;35.

26. Ziyin L, Ueda M. Universal thermodynamic uncertainty relation in nonequilibrium dynamics. Phys Rev Res. 2023;5:013039.

27. Majumdar SN, Bouchaud JP. Optimal time to sell a stock in the Black–Scholes model: comment on ‘Thou shalt buy and hold’, by A. Shiryaev, Z. Xu and X.Y. Zhou. Quantitative Finance. 2008;8(8):753–760.

28. Busseti REK Enzo, Boyd S. Risk-Constrained Kelly Gambling. The Journal of Investing. 2016;25(3):118–134.

29. de Groot DH, Tjalma AJ, Bruggeman FJ, van Nimwegen E. Effective bet-hedging through growth rate dependent stability. Proceedings of the National Academy of Sciences. 2023;120(8):e2211091120.

30. Michel Loreau, Matthieu Barbier, Elise Filotas, Dominique Gravel, Forest Isbell, Steve J. Miller, Jose M. Montoya, Shaopeng Wang, Raphaël Aussenac, Rachel Germain, Patrick L. Thompson, Andrew Gonzalez, and Laura E. Dee. Biodiversity as insurance: from concept to measurement and application. Biological Reviews, 96(5):2333–2354, 2021.

31. Pande J, Fung T, Chisholm R, Shnerb NM. Mean growth rate when rare is not a reliable metric for persistence of species. Ecology Letters. 2020;23(2):274–282.

32. Pande J, Tsubery Y, Shnerb NM. Quantifying invasibility. Ecology Letters. 2022;25(8):1783–1794.

